## Supplementary material for "optoPAD: a closed-loop optogenetics system to study the circuit basis of feeding behaviors"

### Supplementary Table

| figure | line | full genotype | source |
|---|---|---|---|
| 2,3 | *;Gr5a-GAL4;* | *;Gr5a-GAL4; Gr5a-GAL4* | Kristin Scott |
| 2,3,4 | *;;pUAS-Chrimson-mVenus* | *w1118; P{20XUAS-IVS-CsChrimson.mVenus}attP2* | BDSC |
| 2 | *;;57F03-GAL4* | *w1118;; P{GMR57F03-GAL4}attP2* | BDSC |
| 2 | *;;pBDP-GAL4Uw* | *w[1118]; P{y[+t7.7] w[+mC]=GAL4.1Uw}attP2* | BDSC |
| 2,3 | *w[1118];;* | | Barry Dickson |
| 2,3 | *; ;UAS-GtACR1* | *w[1118];; 20xUAS-GtACR1 (attP2)* | Adam Claridge-Chang |
| 3 | *;;Gr-64f-GAL4* | *w[*];; P{w[+mC]=Gr64f-GAL4.9.7}1/TM3, Sb[1]* | BDSC |
| 3 | *;;attP2* | *y1 w67c23; P{CaryP}attP2* | BDSC |
| 3,4 | *Gr66a-GAL4* | *w[*]; P{w[+mC]=Gr66a-GAL4.D}2;* | BDSC |
